# A heterogeneous spatial model in which savanna and forest coexist in a stable equilibrium

**DOI:** 10.1101/400051

**Authors:** Rick Durrett, Ruibo Ma

## Abstract

In work with a variety of co-authors, Staver and Levin have argued that savanna and forest coexist as alternative stable states with discontinuous changes in density of trees at the boundary. Here we formulate a nonhomogeneous spatial model of the competition between forest and savanna. We prove that coexistence occurs for a time that is exponential in the size of the system, and that after an initial transient, boundaries between the alternative equilibria remain stable.

## 1 Introduction

In a 2011 paper published in Science [22], Carla Staver, Sally Archibald and Simon Levin argued that tree cover does not increase continuously with rainfall but rather is constrained to low (*<* 50%, “savanna”) or high (> 75%, “forest”) levels. In follow-up work published in Ecology [23], the American Naturalist [24] and Journal of Mathematical Biology [21], they studied the following ODE for the evolution of the fraction of land covered by grass *G*, saplings *S*, and trees *T*:

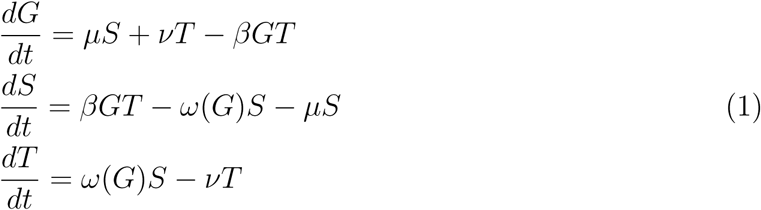

Here *μ ≥ v* are the death rates for saplings and trees, and *ω*(*G*) is the rate at which saplings grow into trees. Fires decrease this rate of progression, and the incidence of fires is an increasing function of the fraction of grass, so *ω*(*G*) is decreasing. Studies suggest (see [24] for references) that regions with tree cover below about 40% burn frequently but fire is rare above this threshold, so they used an *ω* that is close to a step function.

Inspired by this work, Durrett and Zhang [14] considered two stochastic spatial models in which each site can be in state 0, 1, or 2: Krone’s [16] model in which 0 = vacant, 1 = juvenile, and 2 = a mature individual capable of giving birth, and the Staver-Levin foresst model in which 0 = grass, 1 = sapling, and 2 = tree. Theorem 1 in [14] shows that if (0,0,1) is an unstable fixed point of (1) then when the range of interaction is large, there is positive probability of grass and trees surviving starting from a finite set and there is a stationary distribution in which all three types are present. The result they obtain is asymptotically sharp for Krone’s model. However, in the Staver-Levin forest model, if (0, 0, 1) is attracting then there may also be another stable fixed point for the ODE, and as their Theorem 3 shows, in some of these cases there is a nontrivial stationary distribution.

Recently Touboul, Staver, and Levin [26] have investigated a number of modifications of the three species system (1). Variants of the ODE that add a fourth type called forest trees display a wide variety of behaviors including limit cycles, homoclinic, and heteroclinic orbits. Simulations of the spatial version of ODE systems with periodic orbits, see [7], suggest these systems will have stationary distributions that are patchy and with local densities oscillating when the scale of observation is smaller than what physicists call the correlation length, see Figure 4 in [12]. Proving that this occurs is a very difficult problem. Here we will instead focus our attention on a two species simplification studied in [26]

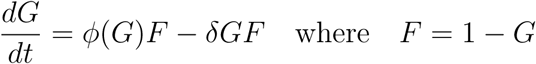

In formulating our version of this model we have two space scales: the dispersal scale *L*, which might be hundreds or thousands of feet, and the continental scale *ML*, which might be thousands of miles. To prove results it is convenient to have the dynamics taking place on a two-dimensional square with a small spacing between points: (ℤ^2^*/L*) *∩* [*−M, M*] ^2^. Tori are convenient mathematically but continents end at oceans so we will use Dirichlet boundary conditions, i.e., the sites outside the square are vacant. The state of site *x* at time *t* can be 0 (grass) or 1 (tree). Each site *x* changes state at a rate dictated by the density of its competitors in a neighborhood of *x*. Motivated by the large variability of rainfall amounts in South America, we will consider a spatially heterogeneous model. Let *f*_*i*_(*x*) be the fraction of type *i* sites in *x* + *𝒩* and let *α >* 1 be an integer. The transition rates are as follows:

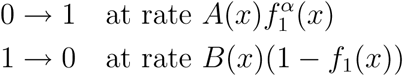

Let 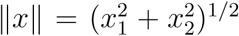 be the Euclidean norm on ℝ^2^, let *D*(*x, r*) = {*y*: *‖y - x‖ ≤ r*} be the disc of radius *r* centered at *x*. To have rotational symmetry in the long range limit we will take *𝒩* = *D*(0, 1), the unit ball. We assume that the climatic conditions vary on a continental scale. To control the variability on intermediate scales, we make the first of several assumptions

(H1) *A*(*x*) = *a*(*x/M*) *and B*(*x*) = *b*(*x/M*) *where a and b are positive and Lipschitz continuous.*

The goal of this study is to show that in our heterogeneous system it is possible to have stable coexistence of grassland and forest, which will not occur in a homogeneous system. Coexistence occurs because we will have regions that are grassland, i.e., mostly 0’s, and regions of forest that are mostly 1’s. The boundaries between the two regions can be predicted from the coefficients *A*(*x*) and *B*(*x*). Specifically, there is a constant *C*_1_ so that {*x*: *A*(*x*)*/B*(*x*) *< C*_1_} is grassland, and {*x*: *A*_1_(*x*)*/B*_1_(*x*) *> C*_1_} is forest. Viewed on the continental scale the boundaries will be stable once equilibrium is reached. Viewed on the dispersal scale, there will be transition zones between the two regions. However, our methods do not tell anything about the nature of these transition zones. Our analysis is done in four steps.

### Step 1

We will let *L → ∞* and scale space by *L* so that the particle system can be replaced by an integro-differential equation. This approach has been used to analyze a number of examples [2, 5, 8, 10, 14, 20, 25] that are not tractable if one assumes a fixed finite range. Before we can state our long-range limit theorem we have to explain what it means for a seqeunce of particle systems *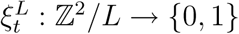* to converge to a function *u*(*t, x*). To do this we pick a small *γ ∈* (0, 1) and tile the plane with squares [*y, y* + *L*^*−γ*^) *×* [*z, z* + *L*^*−γ*^) in such a way that the origin (0,0) is at the lower-left corner of one of the squares. Given an *x ∈*ℝ^2^ let *Q*(*x*) be the square to which *x* belongs and let *u*_*ξ*_(*t, x*) be the fraction of sites *y* in *Q*(*x*) with *ξ*_*t*_(*y*) = 1. We say that 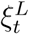 converges to *u*(*t, x*) and write 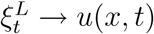 if for any *K* < ∞

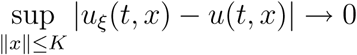

#### Theorem 1

Let 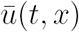 be the average value of u(t, y) over D(x, 1), the ball of radius 1 around x. If 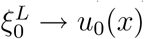 a continuous function then we have 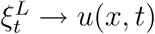 the solution of

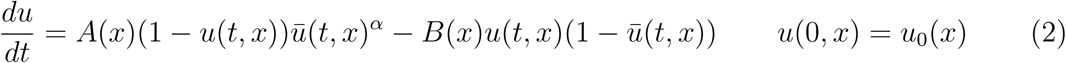

Although it takes some work to prove this, the limit equation is what one should expect from the definition of the process. The site *x* changes 0 *→* 1 at rate *A*(*x*)*f*_1_(*x*)*^α^*, while *x* changes 1 *→* 0 at rate *B*(*x*)(1 *− f*_1_(*x*))*^β^*. The main step in establishing the limit result, is to show that if *L* is large then 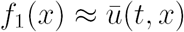.

### Step 2

To analyze the heterogeneous integrodifferential equation (IDE), we will begin by studying the homogeneous ordinary differential equation (ODE)

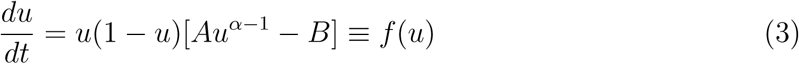

The condition *α >* 1 implies that *f* (*u*) *<* 0 for *u ∈* (0, *ϵ*), so 0 is stable. If *A < B* then *f* (*u*) *<* 0 for *u ∈* (0, 1) so 1 is unstable and there are no interior fixed points. If *A > B* then *f* (*u*) > 0 when *u* is close to 1, so the fixed point at 1 is stable. If *C* = *A/B >* 1 there is an unstable interior at *v* where *Cv*^*α−1*^ = 1, i.e. *v* = *C*^*−1/(α−1)*^.

### Step 3

Durrett and Levin [11] predict that when there is bistability in the limiting ODE (i.e., two stable fixed points) then the density of the equilibrium of the spatial model will be the stronger equilibrium, which can be determined by looking at the sign of the speed of the traveling wave connecting the two fixed points. Since in Step 1 we have taken a limit to get an IDE, we do not need to rely on this heuristic principle. We can use results of Weinberger [27] to prove that this is true for the IDE. That is, there is a constant *C*_1_ so that 0 is the stronger equilibrium and the limit of the IDE if *A/B < C*_1_, and 1 is the stronger equilibrium if *A/B > C*_1_.

### Step 4A

To prove results about the homogeneous system, we use a “block construction.” This technique was invented by in 1988 by Bramson and Durrett [4] and has been used in a large number of examples, see [6]. This application is slightly different than usual since we are on a square and we are proving prolonged survival rather than the existence of a stationary distribution. However, the general outline of the proof is the same.

#### Theorem 2

Let ϵ > 0. (i) Suppose A/B > C_1_. There is a t_0_ < ∞ and an η > 0 so that if L and M are large enough then at times t_0_ ≤ t ≤ exp(ηM ^2^) the fraction of the torus occupied by 1’s is ≥ 1 −ϵ.

(ii) Suppose A/B < C_1_. There is a t_0_ < ∞ and an η > 0 so that if L and M are large enough then at times t_0_ ≤ t ≤ exp(ηM ^2^) the fraction of the torus occupied by 1’s is ≤ E.

This implies in particular that coexistence does not occur in the homogeneous system.

### Step 4B

To describe the behavior of the heterogeneous system now, we divide the plane into regions where *A*(*x*)*/B*(*x*) *< C*_1_ and *A*(*x*)*/B*(*x*) *> C*_1_. To avoid pathologies we assume

(*H*2) *the boundaries A*(*x*)*/B*(*x*) = *C*_1_ *between these regions are given by a collection of smooth curves.*

To prove our result we need another assumption that we later make precise. See the text after Theorem 4

(*H*3) *In each region A*(*x*)*/B*(*x*) *> C*_1_ +*δ there is a sufficiently large patch with a large enough density of 1’s.*

#### Theorem 3

Let δ > 0 and ϵ > 0. There is a time t_0_ < ∞ and η > 0 so that if L and M is large then for t_0_ ≤ t ≤ exp(ηM ^2^)

(i) In each of the regions A(x)/B(x) > C_1_ + δ, the fraction of 1’s is ≤ϵ

(ii) In each of the regions A(x)/B(x) < C_1_ − δ, the fraction of 1’s is ≥ 1 −ϵ

The remainder of the paper is devoted to the proof of this theorem. Steps 1, 3, 4A, and 4B are carried out in Sections 2, 3, and 4. Since this method has been applied to a large number of examples, we describe the ideas involved and refer to previous work for readers want more details.

The analysis in Step 2 can be carried out for the more general model with

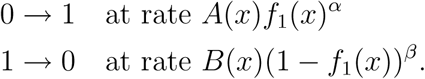

with *α >* 1 *> β >* 0, or even to models with

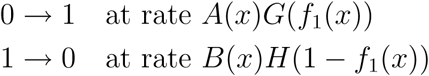

with suitable assumptions on *G* and *H*, which we now state. If *G*^*′*^(0) = 0 and *H*^*′*^(0) = *∞* then *f* (*u*) *<* 0 for small *u* and for *u* close to 1. Let *g*(*y*) = *G*(*y*)*/y* and *h*(*y*) = *H*(*y*)*/y*. If *C* is large enough then

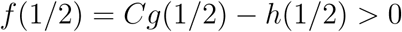

then there are points *v*_1_ *<* 1*/*2 *< v*_2_ with *f* (*v*_*i*_) = 0. 0 and *v*_2_ are stable fixed points and *v*_1_ and 1 are unstable. For our analysis to work, we cannot have more than two interior fixed points. Writing the condition for a fixed point as

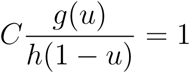

we see that in order to have at most two interior fixed points for all *C* there must be a *w* so that

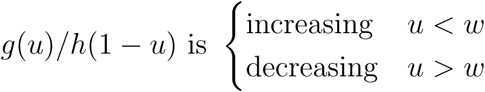

The main obstacle in proving this is that lack of a suitable dual process. In Section 2, we show that when *α* is an integer and *β* = 1, the process can be constructed from a graphical representation in which *α* neighbors are sampled at potential birth events, and one neighbor is sampled at potential death events. For the more general dynamics introduce above we can instead sample *k*_*L*_ *→ ∞* neighbors to obtain an estimate of the local density and let *k*_*L*_ slowly to establish the hydrodynamic limit in Step 1. This approach leads to a number of technical complications so we will not pursue these generalization here.

## 2 Hydrodynamic limit

Since the coefficients change slowly on the dispersal scale, it is enough to consider the homogeneous case *A*(*x*) *≡ A* and *B*(*x*) *≡ B*. For each site *x ∈*ℤ^2^*/L* we have a rate *A* Poisson process 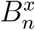 of births and a rate *B* Poisson process 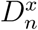 of deaths.

- At each time 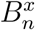 we have *α* random variables 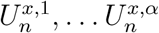 that are drawn from *x* + *𝒩* without replacement. If *x* is vacant at time 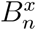 and all the 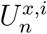 are occupied then *x* becomes occupied.
- At each time 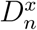 we have a random variables 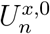 that is uniform on *x* + *𝒩*. If *x* is occupied at time 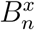 and 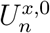 is vacant then *x* becomes vacant.

Given this structure, which is called a graphical representation, we can define a dual process 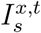 starting at *x* at time *t* and working backwards in order to determine the state of *x* at time *t*. Nothing happens until the first time *s* so that there is a point 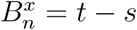 or 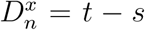. In the first case we add 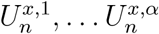 to 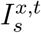. In the second we add 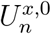. We continue in this way: if there is a 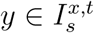 and a point 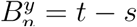 or 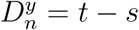 then in the first case we add 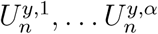 to 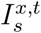. In the second we add 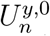. Durrett and Neuhauser [9] call this the *influence set* because if we know the values at 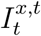 then we can work up the dual process to determine the state of *x* at time *t*.

If at some time, a point that we want to add is already in 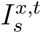 we say that a collision occurs. If *t* is fixed then in the limit as *L → ∞* the probability of a collision tends to 0. In addition

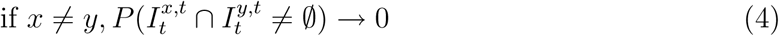

To prove Theorem 1 at this point we can repeat the argument from [2]. The convergence of the dual to a branching random walk implies that the probability that the site *x* is in state 1 at time *t* converges to a limit *v*(*t, x*). Combining (4) with a second moment calculation shows that as *L → ∞* the fraction of occupied sites in a small square [*y*_1_, *y*_1_ + *L*^*−γ*^) *×* [*y*_2_, *y*_2_ + *L*^*−γ*^) converges to *v*(*t, y*), and the fraction of occupied sites in *D*(*x,* 1) converges to 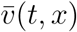, the integral of *v*(*t, x*) over *D*(*x,* 1). With this established, considering the changes of the process over a small interval [*t, t* + *h*] shows that the limit *v*(*t, x*) satisfies the IDE.

## 3 Analysis of the homogeneous IDE

Weinberger [27] studied the asymptotic behavior as *n → ∞* of discrete time iterations *u*_*n+1*_(*x*) = (*Qu*_*n*_)(*x*) where *Q* acts on functions from *u*(*x*): ℝ*^d^ →* [0, 1]. Our evolution occurs in continuous time, but our system can be put into his setting by letting (*Qu*)(*x*) be the solution at time 1, starting from initial condition *u*(*x*). To develop a theory of these iterations Weinberger introduced a number of assumptions. Some of these, like translation invariance, are natural assumptions, but others are somewhat technical. His (3.1), (3.2), (3.3) and (3.5), which are sufficient to obtain the results we will use are satisfied in our case.

In the theory of reaction diffusion equations, traveling wave solutions play an important role. These are solutions to the PDE of the form

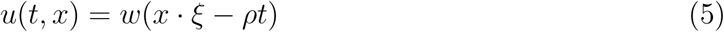

where *ξ* is a fixed direction, i.e., *ξ ∈*𝕊 *^d−1^*, the *d* − 1 dimensional sphere of unit vectors in ℝ*^d^*. This is a plane wave, i.e., at any time *t* the value *u*(*t, x*) on hyperplanes perpendicular to the direction of movement *ξ*. Note that the shape of the wave in (5) does not change and it moves with a constant speed *ρ*.

In the case of a PDE *w* satisfies an ordinary differential equation, but in the case of a discrete iteration or an IDE it is generally not possible to prove the existence of traveling solutions. (For an exception to this rule see [20].) Weinberger shows in      Section 5 of his paper that it is possible to define a wave speed *ρ*(*ξ*) for each direction. In our case the equation is rotationally invariant so all the speeds are the same. Theorems 6.1 and 6.2 in [27] combine to prove the following result in our special case.

Define *C*_1_ so that *ρ >* 0 when *A/B > C*_1_ and *ρ <* 0 when *A/B < C*_1_. Given *C > C*_1_ let *v* = *C*^*−1/(α−1)*^ be the unstable interior fixed point.

### Theorem 4

Suppose ρ > 0 and let 0 < r < ρ < R. (i) If u_0_ = 0 outside a bounded set then

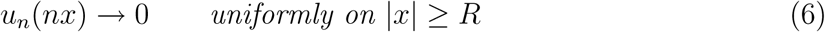

(ii) If σ > v then there is an N_σ_ so that if u_0_ ≥ σ on x_0_ + [0, N_σ_]^2^ then

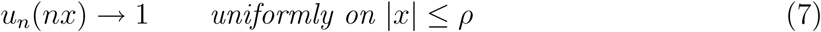

Part (ii) tells us what we need to assume in (H3).

This is a law of large numbers for the solution, i.e., it describes what we see if we scale space by *n* and let *n → ∞*. It does not give information about what is happening near the front. For reaction diffusion equations such as

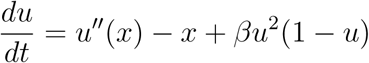

then shape of the solution near the front converges to the traveling wave solution.

The second result will allow us to show that the 1’s survive when *A/B > C*_1_. Things are much simpler when *A/B < C*_1_. Interchanging the roles of 1 and 0 and using Theorem 6.5 of [27] we have:

### Theorem 5

If u_0_ ≤ 1 and u_0_ /≢ 1 then u_n_(x) → 0 uniformly on compact sets as n → ∞.

## 4 Block construction

**Homogeneous case.**Suppose that *A/B > C*_1_. Let *v* be the unstable fixed point of the ODE (3), let *σ > v* and pick *N ≥ N_σ_*, the constant in Theorem 4. Using that result and noting that the distance from (0, 0) to (3*N,* 3*N*) is 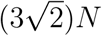 we see that

### Lemma 1

*Let ϵ >* 0. *Suppose the initial configuration of the IDE u*(0, *x*) *≥ σ on* [*− N, N*] ^2^ *and let T* = 10*N/ρ. Part (ii) of Theorem 4 implies that if N is sufficiently large then u*(*T, x*) *≥* 1 *− E on* [*−*3*N,* 3*N*] ^2^.

**Figure 1:**
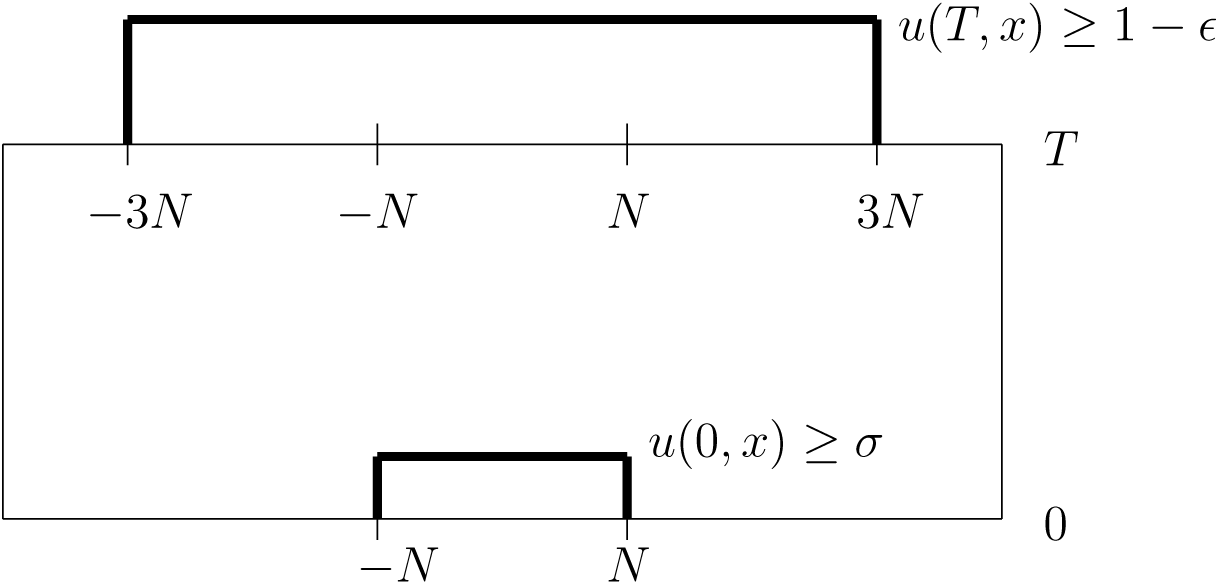
Picture of the block construction in the homogeneous case.

Consider the particle system on ℤ^2^*/L*. If we let *L → ∞*, then by Theorem 1 the particle system converges to the IDE given in (2). As in the formulation of Theorem 1, the statement that particle system *ξ*_*t*_ has density *≥ u*_0_ on a square [*−aN, aN*] ^2^ means that if we tile the plane with squares [*y, y* + *L*^*−γ*^) *×* [*z, z* + *L*^*−γ*^) in such a way that the origin is the lower-left corner of one of the squares, then the density of 1’s in *ξ*_*t*_ in each small square is contained in [*−aN, aN*] ^2^ is *≥ u*_0_.

### Lemma 2

*If the initial configuration of the particle system has density ≥ s* + *ϵ in* [*− N, N*] ^2^ *and N is large then with high probability it will have density ≥* 1 *–* 2*ϵ on* [3*N,* 3*N*] ^2^ *at time T* = 10*N/ρ. In this case we say that the good event G*(0, 0, 0) *occurs.*

If *ϵ* is small 1 2*ϵ s* + *ϵ*. For each (𝓁, *m, n*) ∈ ℤ^3^ with 𝓁 + *m* + *n* even, define a square *I* _𝓁,*m*_ = (2 𝓁*N,* 2*mN*) + [*− N, N*] ^2^. We say that (𝓁, *m, n*) is wet if *ξ*_*nT*_ ≥ 1 *−* 2 *ϵ* on *I* _𝓁,*m*_ at time *nT*. Define the good event *G*(𝓁, *m, n*) by a space and time translation of *G*(0, 0, 0). If (*£, m, n*) is wet and the good event *G*(𝓁, *m, n*) occurs then the four points (𝓁 + 1, *m* + 1, *n* + 1), (𝓁 + 1, *m* + 1, *n* + 1), (𝓁 + 1, *m* + 1, *n* + 1) and (𝓁 *−* 1, *m* + 1, *n* + 1) are wet.

To complete the proof of part (i) of Theorem 2 we will use a comparison with oriented percolation. The first step is show that the good events are *K*-dependent for some fixed value of *K* independent of *N*. To do this we compare the dual with a branching random walk in which (i) at rate *A* a particle at *x* dies and gives birth to particles at *x, U^x,1^*, … *U*^*x,α*^, (ii) at rate *B* a particle at *x* dies and gives birth to particles at *x* and *U*^*x,0*^. To control the movement of the dual we use the following well-known result, see e.g., [3].

### Lemma 3

*Let 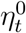 be the particles in the branching random walk at time t starting from a single particle at the origin. If J is chosen large enough, there is a γ >* 0 *so that 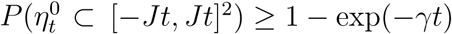.*

Taking *t* = *T* the height of the block in our construction, the form of the error term implies that if *N* is large and *T* = 10*N/ρ* then with high probability 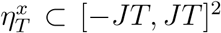 holds for all the starting points in the box [*−*3*N,* 3*N*] ^2^. This implies that the range of dependence between good events on a fixed level *n* is bounded by a *K* independent of *N*. By construction good events on different levels are independent, so we have verified *K*-dependence.

At this point we have shown that, for any fixed *ϵ >* 0, i f *N* i s l a rge e n ough t h en the wet sites in our block construction dominate those in a *K*-dependent oriented percolation in which sites are open with probability *≥*1 *−ϵ*. To prove there is prolonged survival for the oriented percolation process, we could use existing results for the exponential survival time of the two-dimensional oriented percolation, but it is much simpler to turn the two dimensional process on [*− K, K*]^2^ (where *K* = *M/N*) into a one-dimensional process with edges connecting adjacent vertices in the following list:

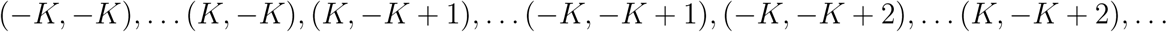

Once we do this it follows from [13] then survival for time exp(*ηM* ^2^) and during that time the density is close that of the stationary distribution of the infinite system, which completes the proof of (i) in Theorem 2. Part (ii) is similar with Theorem 5 is used in place of Theorem 4.

### Heterogeneous case

Consider one component *G* of the open set

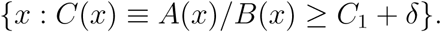

- *A*(*x*) controls the rate at which time is run. We have assumed *A*(*x*) = *a*(*x/M*) where *a* is Lipschitz continuous, so 0 *< α*_0_ *≤ A*(*x*) *≤ α*_1_ *< ∞*.
- If *A* is fixed then the slowest expansion speed *ρ*_0_ occurs when *C*(*x*) = *C*_1_ + *δ*.
- The unstable root is largest when *C* = *C*_1_ + *δ*. Let *v*_0_ = (*C*_1_ + *δ*)*^−1/(α−1)^*

Combining the last three observations and using Theorem 4

### Lemma 4

*Let ϵ >* 0. *Suppose that the initial configuration o f t he I DE u*(0, *x*) *≥ σ on* [*−N, N*] ^2^ *⊂ G let T* = 10*N/ρ*_0_*α*_0_. *If N is large enough then u*(*T, x*) *≥* 1 *− E on* [*−*3*N,* 3*N*] ^2^.

Lemma 4 is the analogue of Lemma 1. It shows that if *u*(0, *x*) *≥ σ* on [*− N, N*] ^2^ then at time *T* we have *u*(*T, x*) *≥* 1 *−ϵ* on [*−*3*N,* 3*N*] ^2^. Combining this result with the hydrodynamic limit in Theorem 1 gives the analogue of Lemma 2 and produces an event that guarantees the expansion of the particle system. Once this is done, we can again compare with oriented percolation. This time we are working in

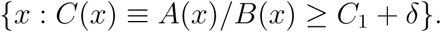

but by using an analogue of the construction for the square we can again compare with a one dimensional process.

## References

[1] Aronson, D. G. and Weinberger, H. F. (1978) Multidimensional nonlinear diffusion arising in population genetics. Adv. in Math. 0 (1978), 33–76

[2] Bessonov, M. and Durrett, R. (2017) Phase transitions for a planar quadratic contact process. Adv. in Appl. Math. 87, 82–107

[3] Biggins, J.D.(1977) Chernoff’s Theorem in the Branching Random Walk J. Appl. Prob. 14, 630–636

[4] Bramson, M. and Durrett, R. (1988) A simple proof of the stability criterion of Gray and Griffeath. Probab. Theory Related Fields. 80, 293–298

[5] Chan, B., Durrett, R., and Lanchier, N. (2009) Coexistence for a multitype contact process with seasons. Ann. Appl. Prob. 19, 1921–1939

[6] Durrett, R. (1995) Ten Lectures on Particle Systems. Pages 97-201 in St. Flour Lecture Notes. Lecture Notes in Math 1608. Springer-Verlag, New York

[7] Durrett, R. (2008) Coexistence in Stochastic Spatial Models. (Wald Lecture Paper). Ann. Appl. Prob. 19, 477–496

[8] Durrett, R., and Lanchier, N. (2008) Coexistence in host pathogen systems. Stoch. Proc. Appl. 118, 1004–1021

[9] Durrett, R., and Neuhauser, C. (1994) Particle systems and reaction diffusion equations. Ann. Prob. 22, 289–333

[10] Durrett, R., and Neuhauser, C. (1997) Coexistence results for some competition models. Ann. Appl. Probab. 7, 10–45

[11] Durrett, R., and Levin, S. (1994) The importance of being discrete (and spatial). Theoret. Pop. Biol. 46, 363–394

[12] Durrett, R., and Levin, L. (1998) Spatial aspects of interspecific competition. Theoret. Pop. Biol. 53, 30–43

[13] Durrett, R., and Liu X-F. (1989) The contact process on a finite set. Annals of Probability 16, 1158–1173

[14] Durrett, R., and Zhang, Y. (2015) Coexistence of Grass, Saplings and Trees in the Staver-Levin Forest Model. Annals of Applied Probability. 25, 104–115

[15] Fife, P.C. and McLeod, J.B. (1977) The approach of solutions of nonlinear diffusion equations to traveling frontsolutions. Arch. Rat. mech. Anal. 65, 335–361

[16] Krone, S. M. (1999). The two-stage contact process. Ann. Appl. Probab. 9, 331–351.

[17] Liggett, T. M. (1985) Interacting particle systems. Springer-Verlag, New York

[18] Mountford, T.S. (1993) A metastable result for the finite multidimensional contact process. Canadian Math Bulletin. 36, 216–222

[19] Mountford, T.S. (1999) Existence of a constant for finite system extinction. J. Statistical Physics. 96, 1331–1341

[20] Neuhauser, C. (1994) A long range sexual reproduction process. Stochastic Process. Appl. 53, 193220.

[21] Schertzer, E., Staver, A. C. and Levin S. (2015). Implications of the spatial dynamics of fire spread for the bistability of savanna and forest. J Math Biol. 70, 329–341

[22] Staver, A. C., Archibald, S., and Levin S. (2011). The global extent and determinants of savanna and forest as alternative biome states. Science 334, 230–232

[23] Staver, A. C., Archibald, S. and Levin S. (2011). Tree cover in sub-Saharan Africa: Rainfall and fire constrain forest and savanna as alternative stable states. Ecology 92, 1063–1072

[24] Staver, A. C. and Levin S. (2012). Integrating theoretical climate and fire effects on savanna and forest systems. Amer. Nat. 180, 211–224

[25] Swindle, G. (1990). A mean field limit of the contact process with large range. Probab. Th. Rel. Fields 85, 261–282.

[26] Touboul, J.D., Staver, A.C. and Levin, S.A. (2018) On the complex dynamics of savanna landscapes. Proc. Natl. Acad. Sci. 115 (7), E1336–E1345

[27] Weinberger, H.F. (1982) Long time behavior of a class of biological models. SIAM J. Math. Anal. 13, 353–396

